# Repeated losses of self-fertility shaped heterozygosity and polyploidy in yeast evolution

**DOI:** 10.1101/2025.09.12.675800

**Authors:** Nina Vittorelli, Cintia Gómez-Muñoz, Irina Andriushchenko, Louis Ollivier, Nicolas Agier, Stéphane Delmas, Yann Corbeau, Guillaume Achaz, Marco Cosentino Lagomarsino, Gianni Liti, Bertrand Llorente, Gilles Fischer

## Abstract

Evolutionary transitions in mating strategy have profound consequences for genetic variation and adaptation. In *Saccharomyces cerevisiae,* mating-type switching is a central feature of the life cycle that enables homothallism, *i.e.,* mating between mitotic descendants of the same haploid cell. Yet heterothallic isolates that have lost this ability are found across diverse niches, indicating that this trait is polymorphic. Here we experimentally characterized loss of mating-type switching in a representative panel of strains. Analysis of 117 telomere-to-telomere genome assemblies revealed multiple independent loss-of-function mutations in the Ho endonuclease gene and structural variants in the silent *HML* and *HMR* cassettes, the three loci essential for switching. We estimated that at least 13 independent transitions from homothallism to heterothallism have occurred in the species history. Analysis of the *HO* genotype of 2,915 strains show that at least 27% are heterothallic. We found that heterothallism is strongly associated with polyploidy and elevated genome-wide heterozygosity, although the strength of these associations varies between populations. Heterothallic isolates are most prevalent in domesticated and clinical clades, consistent with an origin linked to human-associated environments. However, they are also found, though less frequently, in natural niches. Signatures of recombination in *HO* sequences suggest that outcrossing contributed to the ecological and geographical distribution of the trait. Our findings reveal that mating-type switching has undergone repeated losses in *S. cerevisiae* evolution, with major consequences for genome architecture and ecological diversification.

**Significance Statement:** Mating-type switching evolved multiple times in yeasts, enabling haploid selfing, which became a key feature of their life cycles. We show that this trait has been lost repeatedly in the history of *Saccharomyces cerevisiae*, producing heterothallic isolates that cannot switch mating type. Analysis of thousands of genomes reveals that these losses involve multiple mutations in the Ho endonuclease and structural changes in the silent mating-type cassettes. Heterothallism is associated with polyploidy and increased genome-wide heterozygosity. It is more frequent in domesticated and clinical lineages, but also occurs in natural ecological niches. Our findings illustrate how repeated changes in a single life-history trait can reshape genome architecture and highlight the dynamic interplay between genetics, environment, and evolution in a model eukaryote.

## Introduction

Reproductive systems determine how genetic material is passed across generations, thereby shaping the patterns of genetic variation and the processes of adaptation. Reproduction can occur clonally or sexually. In sexual reproduction, the mating system strongly influences the level of inbreeding, *i.e.*, the genetic relatedness between partners. Fungi exhibit a remarkable diversity of reproductive strategies, making them a powerful model group to investigate these questions (1–3). Two paradigmatic mating systems exist in fungi, homothallism and heterothallism. In heterothallic species, mating cannot occur between two mitotic descendants of the same haploid cell. Each haploid cell possesses a mating type determined genetically, and partners need to have different mating types. By contrast, in homothallic species, mating can occur between two mitotic descendants of the same haploid cell, a process called haploid selfing. Different molecular mechanisms can confer homothallism. In some species, haploid cells carry the active genes of both mating types, whereas in others, haploid cells can switch mating type (3). Homothallism maximizes sexual reproductive assurance, ensuring that even a single individual can reproduce, but it does so at the cost of intense inbreeding, which may limit long-term adaptive potential. Heterothallism, on the other hand, fosters genetic diversity and thus enhances adaptability but it exposes the species to sexual reproductive failure when compatible partners are rare. Phylogenomic surveys have revealed that homothallism and mating-type switching have independently evolved on numerous occasions across budding yeasts. Transitions from heterothallism to homothallism occurred far more frequently than the reverse, suggesting an adaptive value to selfing modes under certain ecological conditions (4, 5).

*Saccharomyces cerevisiae* is usually described as a homothallic species capable of haploid selfing through mating-type switching (6). In this species, diploid cells reproduce vegetatively when nutrients are available but undergo meiosis when deprived of nitrogen. Meiosis is coupled with sporulation and produces four haploid spores, which can germinate and mate when nutrients become available again. The mating type of a haploid cell is determined by its *MATa* or *MAT*α allele at the *MAT* locus on chromosome III. Mating can occur only between two cells of opposite mating types, forming a diploid *MATa/MAT*α cell. When no compatible partner is present in the environment of a haploid cell, it can perform haploid selfing by dividing mitotically, then switching mating type and finally mating with its daughter cell (Figure 1A). During mating-type switching, the Ho endonuclease creates a DNA double-strand break (DSB) at *MAT*, which is subsequently repaired by gene conversion with the opposite allele (Figure 1A). Such repair is possible because the sequences of both alleles are stored in two transcriptionally silent loci, the *HMRa* and *HML*α repair cassettes, located in both subtelomeric regions of chromosome III.

**Figure 1.**
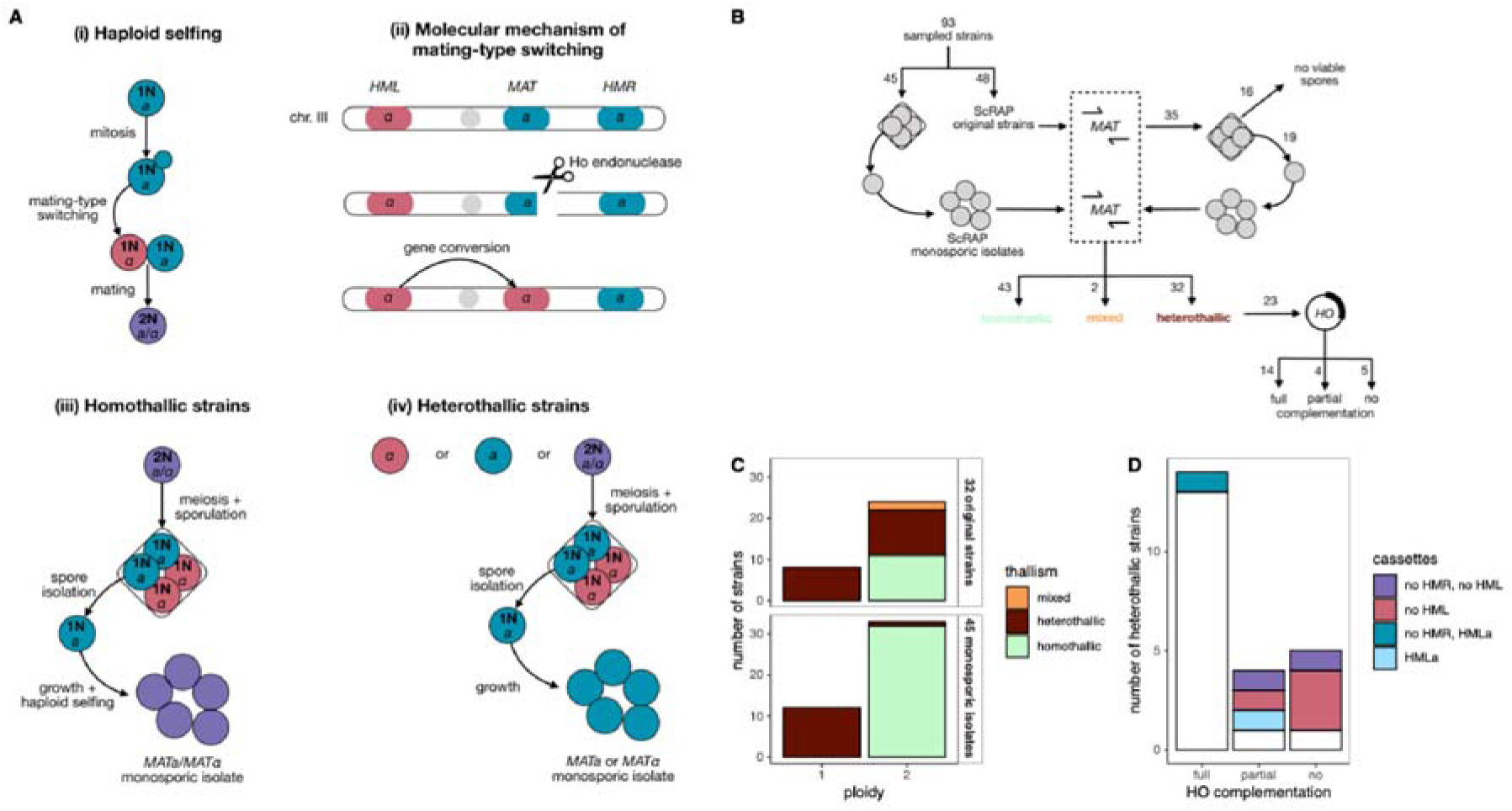
Experimental identification of homothallic and heterothallic strains in the *S. cerevisiae* Reference Assembly Panel (ScRAP, (17)). (A) Haploid selfing through mating type switching (i). During mating-type switching, the Ho endonuclease creates a double strand break in the *MAT* locus, which is repaired by gene conversion using the sequence of the repair cassette *HMR* or *HML* (ii), the grey circle represents the centromere. In homothallic strains, both the original strain and its monosporic isolates carry a *MATa*/*MAT*α genotype (iii) whereas in heterothallic strains, either the original strain or its monosporic isolates carry a single *MATa* or *MAT*α allele (iv). (B) Experimental screen and complementation assay performed in this study. (C) Results of the experimental screen. Monosporic isolates were classified as heterothallic if they carry a single *MATa* or *MAT*α allele, and homothallic if they carry both *MATa* and *MAT*α alleles. Original strains are classified as heterothallic if they or their derived monosporic isolates carried a single *MATa* or *MAT*α allele, and homothallic if they and their derived monosporic isolates carry both *MATa* and *MAT*α alleles. (D) Recovery of homothallism in heterothallic strains after their complementation with a functional *HO* gene. Color indicates unexpected results in the analysis of the *HML* and *HMR* repair cassette (absence, or the mutant *HMLa* allele).

This canonical homothallic life cycle has first been described in a few strains, sampled mainly from vineyards. These strains were *MATa*/*MAT*α diploids, they were sporulated in the laboratory, and so-called monosporic isolates were generated by isolating individual haploid spores. It was observed that these monosporic isolates were also *MATa/MAT*α diploids, indicating that they had switched mating type and that they were therefore homothallic (7) (Figure 1A). However, rare naturally heterothallic strains of *S. cerevisiae* were described very early, notably EM93, isolated in 1938 by Emil Mrak, and its derived lab strains such as S288C (8). Heterothallic strains are characterized by their inability to switch mating-type, and therefore by the maintenance of a *MATa* or *MAT*α mating type. They may be isolated directly with a single *MATa* or *MAT*α mating type. However, most of the time, heterothallic strains are isolated as *MATa*/*MAT*α strains, and they are characterized as heterothallic if their monosporic isolates maintain *MATa* or *MAT*α mating types (Figure 1A). In wine populations, strains were mostly sporulating homothallic diploids with variable levels of genome-wide heterozygosity (7, 9). These observations led to the formulation of the *S. cerevisiae* Genome-Renewal theory, according to which heterozygous mutations accumulated during long periods of vegetative growth and were completely erased by haploid selfing during rare sexual cycles (7, 10). The central role of haploid selfing in *S. cerevisiae*’s life cycle was further supported by inter-species comparative studies, since the *HO* gene and the repair cassettes required for mating-type switching are evolutionary conserved in several related genera, including *Saccharomyces, Nakaseomyces* and *Zygosaccharomyces* (11).

Since then, sampling campaigns have revealed the presence of *S. cerevisiae* all around the world and in multiple ecological niches, from human-associated (*e.g.*, various food, beverage and biofuel productions as well as clinical human samples), to human-remote environments (*e.g.*, primary forest soils, trees, insect guts)(12–16). Genomic surveys uncovered a complex population structure with a great diversity in genetic patterns, including ploidies from 1N to 5N, aneuploidies, structural variants and various levels of genome-wide heterozygosity (13, 17). This suggested a previously underestimated variability of the life cycle within the species (18). It has then been shown that in some domesticated populations, the ability to enter the sexual cycle is reduced, with lower levels of both sporulation and spore viability (19). Additional heterothallic isolates have been described in population-specific studies: in a clinical sampling (20), in a local wild site in Mount Carmel in Israel (21) and in Norwegian breweries (22). In these studies, heterothallism was always associated to the presence of loss-of-function (LoF) mutations in the coding sequence of the *HO* gene, preventing mating-type switching (21–23). Here, after having experimentally confirmed the frequent occurrence of heterothallic isolates in the *S. cerevisiae* population, we examined how often transitions from homothallism to heterothallism occurred in *S. cerevisiae*, investigated their molecular basis, reconstructed their ecological and evolutionary history, and assessed their consequences for patterns of genetic variation at the species level.

## Results

### *HO* inactivation and silent cassette alterations cause frequent heterothallism

We first examined the prevalence of heterothallism using the *S. cerevisiae* Reference Assembly Panel (ScRAP, (17)), a collection of strains with telomere-to-telomere (T2T) and haplotype-resolved genome assemblies (Dataset S1, Figure S1). These assemblies provide access to phased alleles in heterozygous contexts and to gene content within complex subtelomeric regions, which was essential for assessing the functional consequences of sequence variations at the *HO*, *HML*, and *HMR* loci.

We experimentally screened 93 haploid and diploid ScRAP strains without artificial *HO* deletion for mating-type switching ability, *i.e.*, thallism (Dataset S2, Supporting Information). We identified 32 homothallic and 43 heterothallic strains as well as two strains with mixed phenotypes. The remaining 16 strains did not produce viable spores, so their selfing ability could not be assessed at this step (Figure 1B). More specifically, we genotyped the mating type locus of the 45 monosporic isolates that were generated and sequenced in the frame of the ScRAP study (17). We identified 32 homothallic diploids which carried a dual *MATa*/*MAT*α genotype, as well as 13 heterothallic strains which maintained a single mating type (five *MATa* and eight *MAT*α) (Figure 1C). All heterothallic strains were haploid, except one *MAT*α diploid, possibly corresponding to a case of endoreduplication (24). We also genotyped the mating type of the remaining 48 ScRAP original strains (Figure 1C). We found 13 heterothallic isolates with a single mating type (eight *MATa* and five *MAT*α), including eight haploids and five diploids, possibly corresponding to cases of endoreduplication (24). We sporulated the 35 remaining *MATa*/*MAT*α diploids to derive monosporic isolates. We identified 16 diploid strains with defective sporulation or unviable spores. We genotyped the monosporic isolates from the 19 remaining diploid strains and found 6 heterothallic strains and 11 homothallic strains. The monosporic isolates of the last two strains showed mixed phenotypes: CFF produced *MATa* (10/31) and *MATa/MAT*α (21/31) monosporic isolates, while UWOPS034614 produced *MATa* (3/12), *MAT*α (3/12), and *MATa/MAT*α (6/12) monosporic isolates.

We tested whether heterothallism resulted from the inactivation of the Ho endonuclease function by performing a complementation assay. We transformed 23 ScRAP heterothallic strains with the pHS2 plasmid carrying a functional *HO* gene under the control of its native promoter, and then genotyped the *MAT* locus of two transformants per strain to assess restoration of the mating-type switching ability. Among the 23 strains, 14 exhibited full *HO* complementation, four partial complementation and five no complementation (Figure 1D). More specifically, we first complemented 18 single-mating-type isolates and obtained *MATa*/*MAT*α genotypes in 11 cases (full *HO* complementation), the mating type of the initial strain in five cases (no *HO* complementation), whereas in the two remaining cases (*MAT*α strains), both transformant clones had switched to *MATa* (partial *HO* complementation), indicating a partial restoration of the switching ability, but with additional factors causing the inability to restore the *MATa*/*MAT*α genotype (Figure S2A). We then tested five diploid strains (*MATa*/*MAT*α) by transforming at least four of their single-mating type monosporic isolates. In three cases, all transformants from all monosporic isolates were *MATa*/*MAT*α (full *HO* complementation), whereas partial complementation was observed in the two others (Figure S2B). In one case (BAF), the four monosporic isolates from the same tetrad were analyzed and while all transformants recovered the *MATa*/*MAT*α for two monosporic isolates (one *MATa* and one *MAT*α), all transformants were *MATa* for the two other *MATa* and *MAT*α monosporic isolates. In the last case (ASN), five monosporic isolates from three tetrads were transformed and partial *HO* complementation was observed in two of them (Figure S2B). Overall, among the 23 heterothallic strains that were tested, at least 18 exhibited full or partial *HO* complementation, confirming that the absence of a functional Ho protein is the main cause of heterothallism. However, additional factors also contribute to heterothallism in at least nine isolates, five of which show no *HO* complementation and four show partial complementation.

We analyzed the T2T phased genome assemblies of the 117 haploid, diploid and polyploid ScRAP strains devoid of artificial *HO* deletion (including the 93 strains used for the experimental screen; Supporting Information), to identify potential sequence alterations in the *HML* and *HMR* repair cassettes. All retrieved *HMR* loci carried the expected *MATa* sequence (*HMRa*), whereas 12 *HML* loci carried *MATa* (*HMLa*) instead of the expected *MAT*α sequence (*HML*α). These 12 *HMLa* cassettes were found in the genome assembly of nine isolates (Dataset S1), among which four were heterothallic isolates, one isolate showed both homothallic and heterothallic phenotypes (CFF) and four isolates were not experimentally characterized (one non-sporulating diploid and three tetraploids). Such strains that carry a mutant *HMLa* allele lack the silent α-donor sequence and are therefore unable to switch from *MATa* to *MAT*α. This allelic defect explains asymmetric mating type switching in CFF as well as partial *HO* complementation in BAF (Figure 1D). These findings show that rearrangement in the *HML* cassette may lead to partial heterothallism.

We also identified several strains whose T2T genome assemblies completely lacked the *HML* or *HMR* loci (Supporting Information). Although the absence of a cassette in a genome assembly could reflect an assembly artifact, among the five strains showing no *HO* complementation, three lacked *HML* and one lacked both *HML* and *HMR* (Figure 1D). Likewise, two of the four strains with partial *HO* complementation lacked *HML* (Figure 1D). If these cassettes are truly absent, this would explain why these strains are partially or totally unable to switch mating type.

Together, these results demonstrate that heterothallism in *S. cerevisiae* arises primarily from Ho inactivation, with additional contributions from structural alterations or loss of the silent *HML* and *HMR* cassettes, highlighting multiple molecular routes that disrupt mating-type switching.

### Life cycle transitioned to heterothallism at least 13 independent times

We extracted 151 full *HO* coding sequences from the T2T genome assemblies of the 117 ScRAP strains. We identified 160 variants compared to the functional reference sequence, including 67 synonymous single nucleotide variants (SNVs), 81 missense SNVs, four nonsense SNVs (236T>A, 238C>T, 1153G>T, 1731T>A), five frameshift deletions, one frameshift insertion, a loss of start codon (3G>A) and one in-frame 108 bp deletion (1564- 1671, previously observed in (25, 26)) resulting in a 36 amino-acid deletion (Figure 2A, Dataset S3). We combined our results from the experimental screen and the complementation assay with previously published data to classify all variants according to their predicted effect on Ho protein functionality. We first categorized as neutral 90 variants that include the 67 synonymous variants, 20 non-synonymous variants present in the sequence of homothallic strains, and 3 variants previously described as neutral (565A>G, 1214T>C and 1424A>T) (23) (Figure 2A, Figure S3). We then defined as loss-of-function (LoF) 18 variants consisting of the four nonsense mutations, the start codon loss, six frameshifts and seven missense SNVs at functionally critical positions (27), including the 667G>A variant previously shown to inactivate Ho in the reference strain S288C (23) (Figure 2A). The third category comprised all variants for which the effects on Ho functionality remain unknown. Overall, out the 160 *HO* variants from the 151 *HO* coding sequences of the ScRAP dataset, 56% are predicted to be neutral regarding protein functionality, and at least 11% likely abolish protein function.

**Figure 2.**
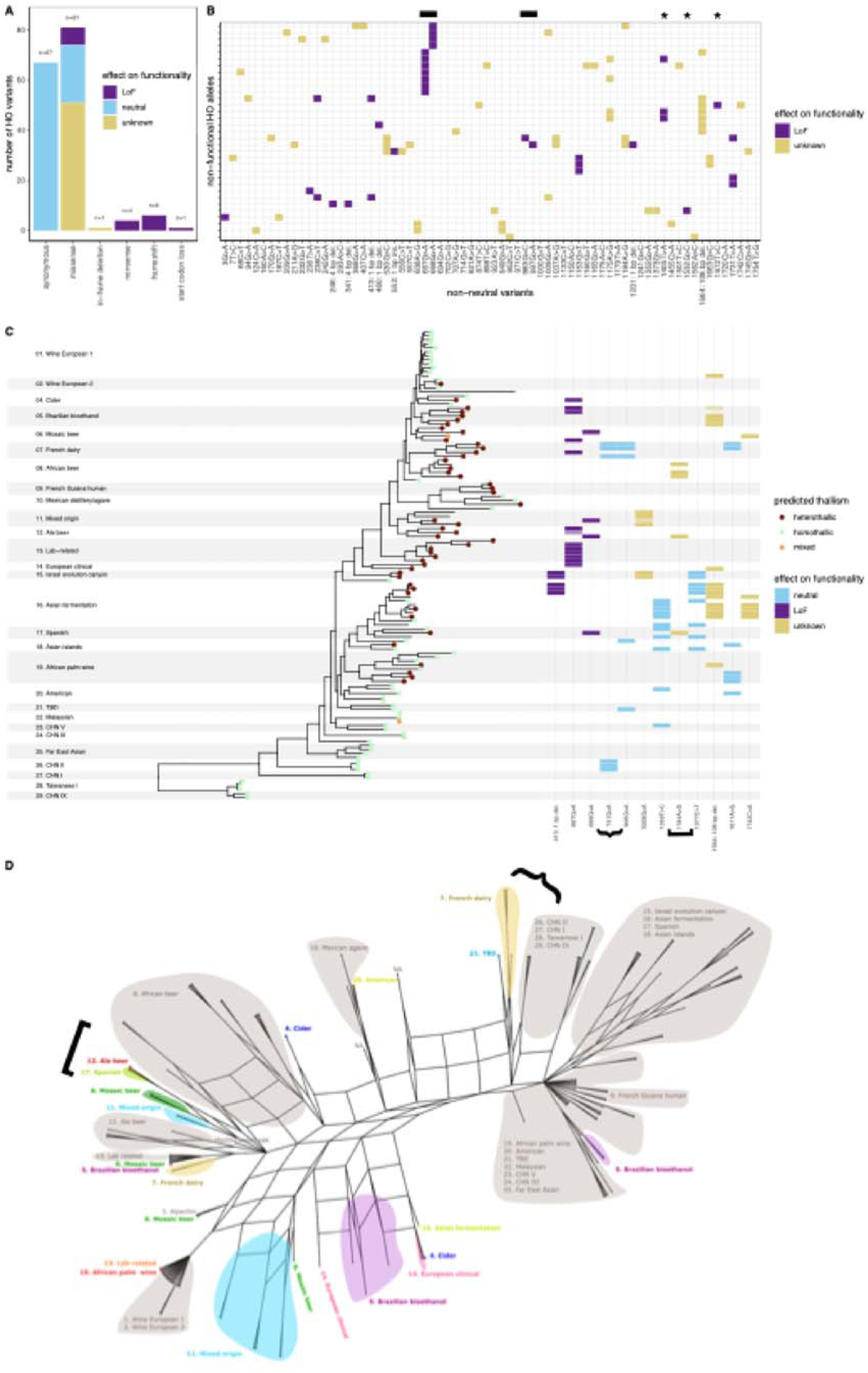
Functional and evolutionary analysis of the *HO* gene in the ScRAP. (A) Types of variants in the *HO* coding sequences and predicted effects on protein functionality. (B) Distribution of non-neutral variants in non-functional *HO* alleles. The x-axis shows the 70 non-neutral variants (18 LoF variants + 52 variants with unknown functionality), and the y-axis shows the 33 non-functional alleles. Black rectangles at the top indicate functionally critical positions in the two LAGLIDADG domains required for Ho endonuclease activity, while black stars indicate functionally critical positions in the three zinc-finger domains involved in DNA binding (27). (C) Phylogenetic distribution of homo- and heterothallic strains and selected *HO* variants. *Left:* the whole-genome-based tree and the clade names are from (17). *Right*: lighter colored rectangles indicates that the variant is present at the heterozygous state. (D) *HO* sequence network. Grey areas indicate clades whose phylogenetic relationships match the whole-genome tree, while colored areas and dots highlight clades with split clustering, where individual strains are positioned far from their cognate clades. In panel C and D, the curly bracket indicates that the proximity of the French dairy and CHN II strains in the network is consistent with the phylogenetic distribution of the 741G>A variant; while the square bracket indicates that the clustering of Ale beer, African Beer and Spanish strains is consistent with the phylogenetic distribution of the 1184A>G variant.

We identified 80 different *HO* alleles (*i.e.*, unique combinations of variants) among the 151 *HO* sequences. We considered as functional the 32 alleles that carried only neutral variants and as non-functional the 33 alleles that carried at least one LoF variant or that were characterized as such in the complementation assay (Figure S4). Additionally, we used these results on *HO* allele functionality to infer the phenotype of the 40 ScRAP isolates that were not testable with the experimental screen (Figure S4). Among them, 16 carried only non-functional *HO* alleles and were therefore predicted heterothallic, 14 carried only functional *HO* alleles and were predicted homothallic, while no prediction could be made for the remaining 10 strains. Overall, by combining the results of the experimental screen and the prediction based on *HO* allele functionality, we estimated that 48% (57/117) of ScRAP isolates are homothallic, 41% (48/117) are heterothallic, 2% (2/117) exhibit both phenotypes, and 8% (10/117) have an unknown phenotype.

We analyzed the distribution of the 70 non-neutral variants – comprising the 18 LoF variants and the 52 variants with unknown effect – in the 33 non-functional *HO* alleles. Each of the 70 non-neutral variants occurs in seven or fewer non-functional alleles (Figure 2B, Figure S5). So, it is likely that there were multiple independent origins to these non-functional *HO* alleles. We computed the minimal number of non-neutral mutations required for all non-functional alleles of our dataset to carry at least one such variant. This analysis indicated a minimum of 13 non-neutral mutations. In the absence of recombination, this represents the minimal number of independent loss-of-function events in the *HO* gene, *i.e.,* transitions from a functional to a non-functional *HO* allele. This is a reasonable approximation for the number of transitions from a homothallic to a heterothallic life cycle. However, this is likely an underestimate as some lineages may have transitioned to heterothallism through mutations not captured in our variant set, either within the *HO* coding sequence or elsewhere in the genome. Interestingly, heterothallic strains are absent from the Asian and American clades that diverged early from the species’ last common ancestor (clades 20–29 in Figure 2C), but they are widely distributed throughout the rest of the ScRAP phylogenetic tree. Some clades consist entirely of heterothallic isolates, while many others, including Wine European 2, Cider, Mexican Agave, Asian Fermentation, Spanish, Asian Islands, and African Palm Wine, contain both homo- and heterothallic strains. This pattern suggests either independent losses of switching ability within these clades or recombination between different alleles.

### Recombination is a major driver of *HO* evolution

Recombination events may have contributed to some transitions between homothallism and heterothallism by transferring LoF variants into initially homothallic backgrounds. To investigate this, we analyzed the distribution of variant alleles in the *HO* coding sequences and detected strong signatures of recombination in multiple pairs of biallelic sites where the four possible haplotype combinations were observed across the 151 sequences. For instance, neutral variants 1059T>C and 1377C>T co-occur in four strains, appear individually in seven and five strains respectively, and absent from the remaining 101 strains (Figure 2C). Additionally, several variants show patchy distributions in the population, further supporting a possible role of recombination in *HO* evolution. For instance, the neutral variant 741G>A is found only in the two divergent CHNII and French Dairy populations while the LoF variants 667G>A occurs in some isolates of the Cider, Brazilian Bioethanol, Mosaic Beer, French Dairy, Ale Beer, Lab-related and European Clinical groups (Figure 2C). The 108 bp deletion is present in some but not all isolates of the African Palm Wine, Asian Fermentation, Brazilian Bioethanol, Wine European and European Clinical groups (Figure 2C). Its patchy distribution could reflect recombination, but we cannot exclude recurrent mutational events as the deletion lies between two identical 8bp repeats (TGTATAAA), which may predispose this region to repeated losses.

Given these punctuated signatures of recombination, we asked whether the set of full-length *HO* sequences supported a network-like phylogenetic signal or a simpler tree-like pattern of descent in which recombination plays only a minor role. Phylogenetic reconstruction produced an extensively reticulated network, consistent with multiple recombination events shaping *HO* evolutionary history (Figure 2D). Nonetheless, a strong phylogenetic signal remains visible, with many clades retaining stable clustering compatible with only a moderate impact of recombination (e.g., Wine European 1 and 2, or clades between Israel Evolution Canyon (#15) and CHN IX (#29), Figure 2D). By contrast, other clades revealed a more complex history, with groups split into subclusters (e.g., Brazilian bioethanol (#5) or French dairy (#7)) or with individual strains located far from their cognate clades. These patterns suggest close relatedness of *HO* sequences from isolates that are otherwise genetically distant. For example, some French dairy (#7) isolates cluster next to CHN II (#26), an association also supported by the 741G>A variant found only in these two clades (Figure 2C). Similarly, one Ale beer strain harbors an *HO* sequence closely related to both the African beer (#8) and Spanish (#17) clades, a proximity corroborated by the 1184A>G variant (Figure 2C). These examples further support recombination as a major driver of *HO* sequence diversity.

### At least 27% strains are heterothallic at the species level

Functional predictions of *HO* variants were then extended to the 3,034 Genomes Panel (3,034GP (28)), the largest available collection of *S. cerevisiae* Illumina genome sequences, which includes both a catalog of nucleotide variants (SNPs) and the information on the heterozygosity and ploidy levels for 3,039 strains. This broader dataset allowed us to infer the species-wide prevalence of heterothallism and to evaluate its association with heterozygosity and polyploidy across clades and ecological niches.

We inferred the homo- or heterothallic phenotype of the 3,034GP strains based on their *HO* genotypes. To focus exclusively on naturally occurring heterothallism, we excluded 124 strains with an artificial *HO* deletion (13). We retrieved the *HO* genotypes of the remaining 2,915 strains and identified 395 variants relative to the reference functional sequence. Using the same classification scheme as for the ScRAP sequences, 10% (40/395) of variants were predicted to cause LoF, 37% (146/395) were classified as neutral for the Ho protein functionality and 53% (209/395) had an unknown effect on Ho protein functionality (Figure 3A, Dataset S4). Most of the variants were rare, with 71% being present in 10 strains or less (282/395, Figure 3B). The three most frequent LoF variants – 668G>A in 289 strains, 667G>A in 249 strains and 646C>T in 153 strains – were all missense variants located in the first LAGLIDADG motif, which involved in the endonuclease activity of the Ho protein (27) (Figure 3C). We found no variant between positions 1560 and 1681, which correspond to the 108bp deletion identified in the ScRAP sequences, nor between positions 1708 and 1761 (Figure 3C). A majority of variants were present at heterozygous state in at least one strain (247/395, *i.e.*, 63%), and sometimes in multiple strains (Figure S6). For instance, the most frequent LoF variant, 668G>A, occurs in 62 strains in a homozygous state, and in 227 strains in a heterozygous state.

**Figure 3.**
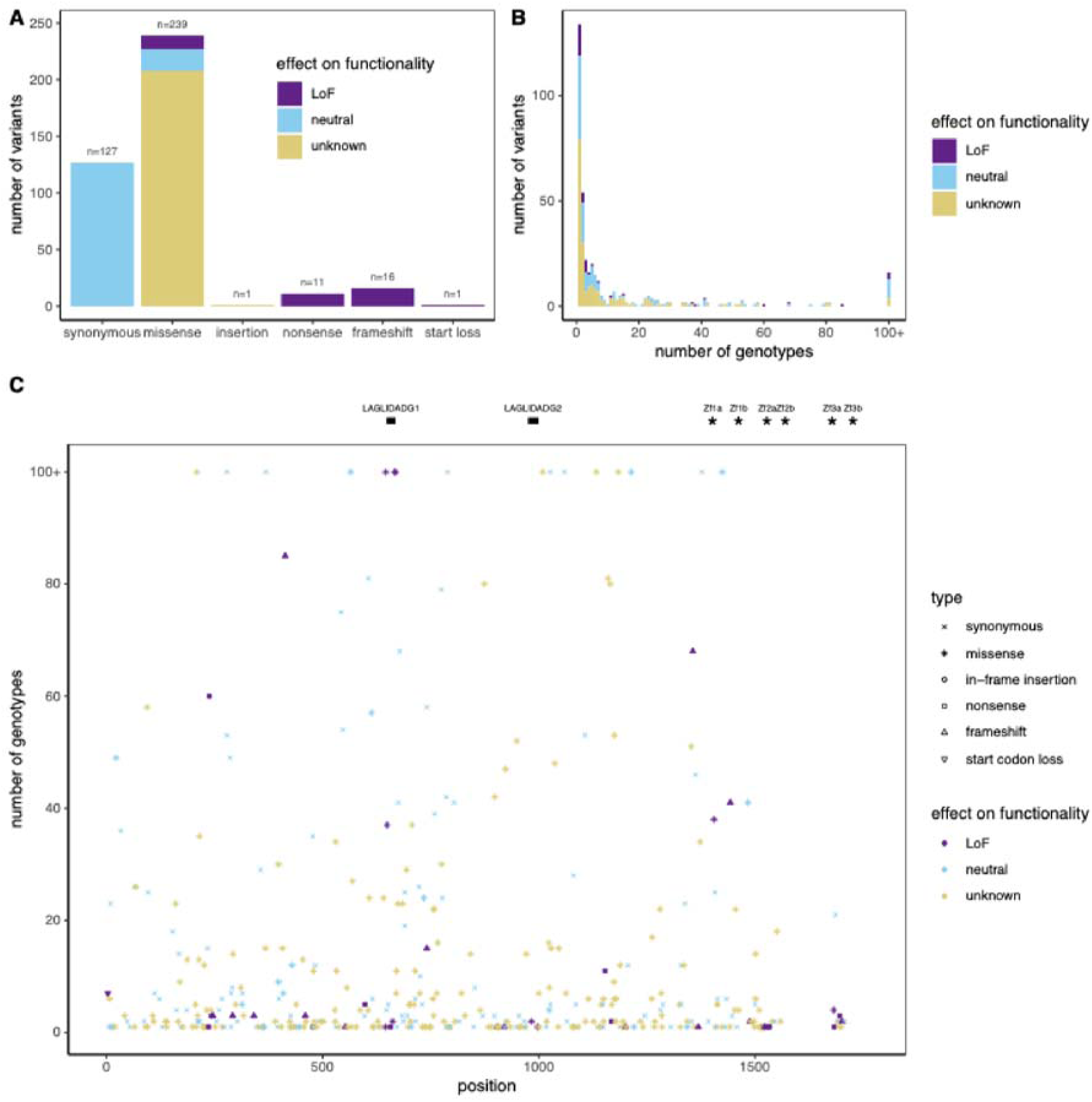
*HO* coding sequence variants in the 3,034 Genome’s Panel (3,034GP (28)). (A) Types of variants found in the *HO* genotypes of the 2,915 strains without artificial *HO* deletion from the 3,034GP. (B) Number of genotypes in which the variant is found. (C) Position of the variants in the 1761 bp *HO* coding sequence. At the top, black rectangles indicate the position of the two LAGLIDADG motifs involved in the endonuclease activity; whereas black stars indicate the position of the three pairs of zinc-finger motifs, all critical for DNA binding except Zf3b (27).

We used this variants classification to predict the switching ability of the 2,915 strains, considering functional *H0* alleles as a proxy for homothallism, and non-functional *HO* alleles as a proxy for heterothallism. A total of 1,685 strains (58%) contained only neutral variants and were classified as homothallic. Menwhile, 779 strains (27%) contained at least one LoF variant and were classified as heterothallic, among which 456 were fully heterothallic (at least one LoF variant in a homozygous state) and 323 were at least partly heterothallic (all LoF variants in a heterozygous state, meaning that some alleles could still be functional depending on the segregation of the LoF variants). The remaining 451 strains (15%) have undetermined thallism status (Dataset S5). We likely underestimate the proportion of heterothallic since we may have missed LoF variants, either because they had unknown effect on *HO* functionality or because they reside in other genes involved in mating-type switching. Overall, we estimate that heterothallic strains represent at least 27% of the 3,034GP collection.

### Heterothallism is associated with polyploidy and genome-wide heterozygosity

We assessed the association between heterothallism, genome-wide zygosity and polyploidy. Strains containing more than 500 heterozygous SNPs genome-wide were considered heterozygous whereas the rest were considered homozygous (28). Ploidy was measured by flow cytometry for 1,278 strains while for the remaining strains, ploidy could be inferred only in heterozygous strains based on their allele balance ratio (28) (Figure S7).

Among the 488 polyploid strains from the 3,034GP collection, we identified a majority of heterothallic strains (358, *i.e.*, 73%) and only 75 homothallic strains (15%) (Figure 4A). On the opposite, among the 1,765 diploids, we identified a majority of homothallic strains (1,056, *i.e.*, 60%) and only 374 heterothallic strains (21%) (Figure 4A). However, heterothallic strains tend to be as often diploid (48%, 374/779) as polyploid (46%, 358/779). The strong association between heterothallism and polyploidy suggests a potential causal link, although its direction cannot be conclusively determined. Additionally, heterothallic strains are almost all heterozygous (91%, 712/779), and very rarely homozygous (8%, 59/779), suggesting that heterothallism promotes heterozygosity, likely because it prevents haploid selfing (Figure 4B). However, among the 1,749 heterozygous strains, we found similar proportions of heterothallic (712/1749, 41%) and homothallic strains (673/1749, 38%). On the contrary, among the 1,152 homozygous strains, we find only 5% heterothallic strains (59/1152), and 88% homothallic strains (673/1749, 38%). These results suggest that the heterothallic life cycle strongly promotes heterozygosity, but that is not the only reproductive regime to generate heterozygosity.

**Figure 4.**
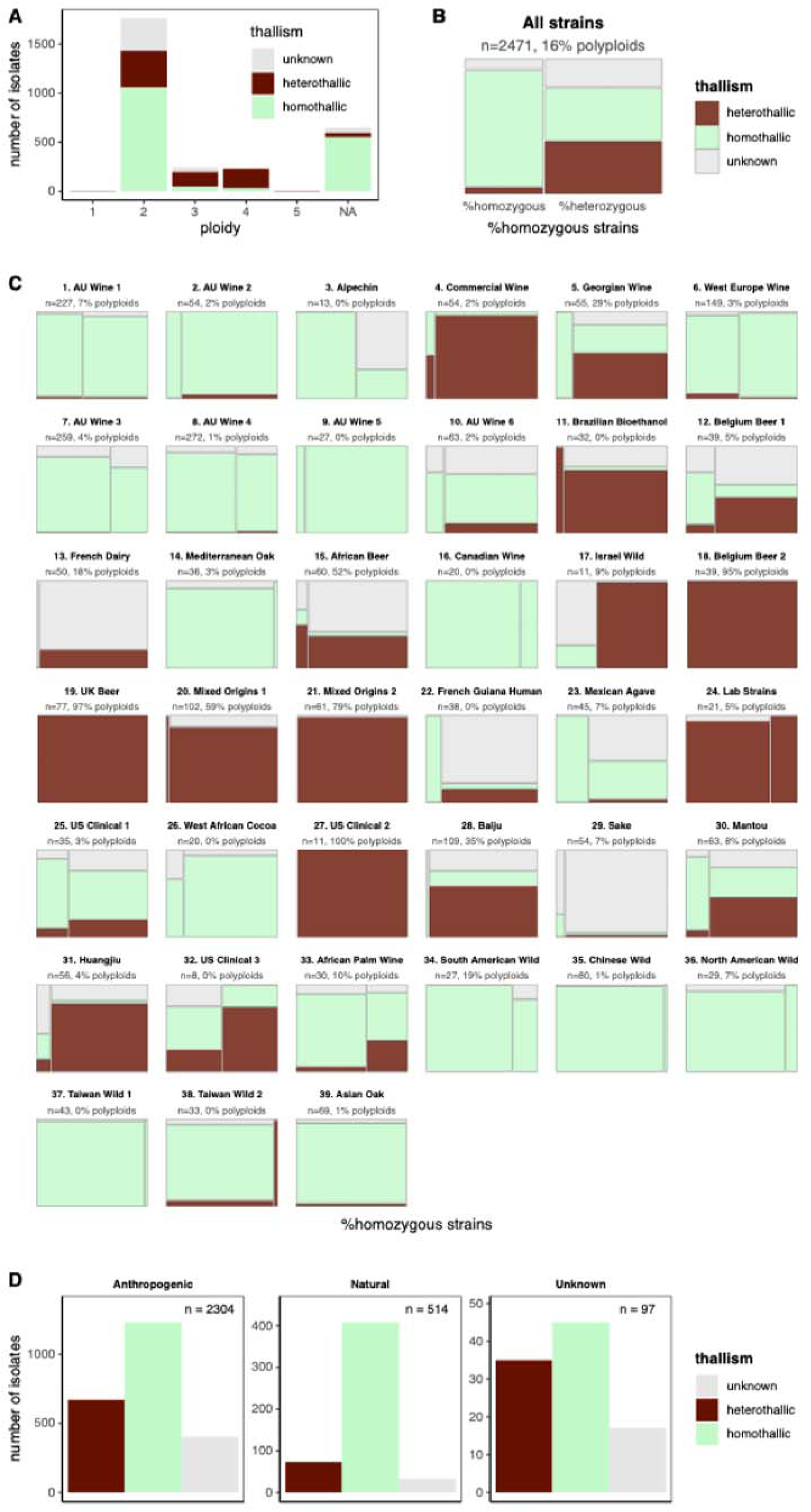
Association between thallism, ploidy, heterozygosity and ecology. (A) Distribution of thallism and ploidy across 2,915 strains. (B) Thallism and genome-wide zygosity of 2,471 strains. The 444 strains with undefined genome-wide zygosity or unattributed clades were excluded. The area of each rectangle represents the proportion of strains for each combination of genome-wide zygosity (x-axis) and thallism (color). (C) Thallism and genome-wide zygosity by clade. The set of strains and the representation are as in panel B. All strains of clade 39 are homozygous, all strains of clades 18, 19, 21, 27 are heterozygous. (D) Thallism across 2,915 strains by ecological category.

Overall, our results show that heterothallism is strongly associated with polyploidy and genome-wide heterozygosity at the species scale. However, there are heterothallic diploid strains, as well as homothallic heterozygous strains. This suggests that multiple interactions exist between thallism, ploidy and zygosity. We investigated if we found different combinations of these traits within the different populations of the species.

### Contrasting reproductive strategies shape genomic diversity across ecological niches

We investigated the prevalence of heterothallic strains within the different clades of the *S. cerevisiae* species (Figure 4C). Several domesticated and clinical clades contain mostly heterothallic strains (>80%), among which Commercial Wine, Belgium Beer 2, UK Beer, Mixed Origins 1 and 2, Lab strains, US Clinical 2. All these clades mainly comprise heterozygous strains (>90%) except for the Lab strains. By contrast, the proportion of polyploids varies widely, ranging from 2% in Commercial Wine, to 59% in Mixed Origins 1, 79% in Mixed Origins 2 and exceeding 95% in Belgium Beer 2, UK Beer, and US Clinical 2. These patterns suggest that the link between heterothallism and polyploidy is population-dependent. Polyploidy may be more counter-selected in populations with frequent sexual cycle, since it impairs meiosis. Among the clades that contain mostly homothallic strains (>80%), we found wine, wild and West African Cocoa clades. All these clades have very low proportions of polyploids, except for the South-American Wild Clade which contains 19% polyploids. We observe that most wild clades contain mainly homozygous strains (>85%), whereas wine clades and the West African Cocoa clade contains higher proportions of heterozygous isolates (from 15% to 93%). This suggests that, despite sharing a homothallic life cycle, wild and wine populations differ in reproductive behaviors. In wine populations, higher heterozygosity levels may be promoted by longer vegetative phases or more frequent outcrossing. Finally, we found four clades with both homothallic and heterothallic strains (>25% each): Georgian Wine, Belgium Beer 1, Mantou et US Clinical 3. Among these, we find variable proportions of heterozygous isolates (50% to 85%), and low proportions of polyploids (<8%) except in the Georgian Wine clade (29% polyploids). Overall, heterothallic isolates were found primarily in clinical and domesticated clades, with the exception of wine, alpechin and Mexican agave clades, while homothallism dominates wild and wine clades. Interestingly, some populations maintain both behaviors, demonstrating that multiple mating strategies can coexist in balanced proportions within the same population.

Using sampling information, we classified strains by ecological niches independently of their clade of origin. Niches were categorized as anthropogenic (associated with human activity), natural (associated with wild plants, animals, or environmental samples), or unknown (Dataset S5). We then compared the distributions of heterothallic and homothallic isolates across these ecological categories (Figure 3D). Among the 2,304 strains isolated from anthropogenic niches, heterothallism was relatively common with 29% of strains heterothallic (671/2322), 53% homothallic (1231/2322) and 17% of unknown thallism (403/2322). By contrast, strains from natural niches showed a lower but still notable presence of heterothallism with 14% heterothallic (73/514), 79% homothallic (408/517) and 6% of unknown thallism (33/517). Among the 73 heterothallic strains that were isolates in a natural niche, 24 came from the Mixed Origins 1 clade, seven from the Israel Wild clade (as already described in (21)) and six from the Huangjiu clade. These findings indicate that while heterothallic isolates are overrepresented in human-associated environments, some heterothallic strains can persist in natural niches, either after originating from anthropogenic environments and returning through feralization, or through the independent loss of homothallism in nature.

## Discussion

Overall, our results reveal a that heterothallism in *S. cerevisiae* is more prevalent than usually thought, with at least 27% heterothallic strains in the 3,034GP, the most comprehensive set of strains studied to date. We found multiple independent transitions from homothallism to heterothallism in *S. cerevisiae*. In populations where mating-type switching and haploid selfing occur frequently, LoF mutations in *HO* likely arise but are eliminated by negative selection. This interpretation is consistent with Mortimer Genome-Renewal theory (10) which proposes that haploid selfing enables efficient purifying selection, *i.e.* selection against deleterious mutations in environments to which the organism is already well adapted. By contrast, in ecological niches more distant from the ancestral conditions, or in changing environments, two non-mutually exclusive hypotheses could explain the persistence of heterothallic strains. First, heterothallism may become less costly. Heterothallism is costly if sexual cycles are frequent and compatible partners are scarce. But if sexual cycles are rare (*e.g*., under nutrient rich conditions) or if compatible partners are abundant, selection on mating-type switching may be relaxed. Second, heterothallism may provide benefits by promoting crosses with genetically distant partners. Such outcrossing could be particularly advantageous in some ecological niches where mating between divergent genotypes accelerates the formation of more adaptive combinations. The repeated losses of mating-type switching in *S. cerevisiae* provide a striking example of how key life cycle traits can be modified multiple times within a single species. Such transitions may reflect ecological heterogeneity and the balance between the costs and benefits of selfing versus outcrossing. Our results indicate that these repeated losses primarily result from the dynamic evolution of the *HO* sequence, through the recurrent accumulation of loss-of-function mutations and diversification via recombination. In addition, natural variation in the silent cassettes may also contribute to the emergence of heterothallism, highlighting multiple genetic routes toward the loss of mating-type switching.

We showed that heterothallism is associated with genome-wide heterozygosity. This pattern is expected, since a single round of haploid selfing generates a completely homozygous diploid. In populations with frequent sexual cycles, increased genome-wide heterozygosity in heterothallic strains could reflect a more outcrossed mating system. By contrast, in populations with rare or absent sexual cycles, heterozygosity would arise from the accumulation of mutations during extended vegetative growth, while relaxed selection on sexual mechanisms including mating-type switching would promote heterothallism. These two explanations for the association between heterothallism and genome-wide heterozygosity are not mutually exclusive, and their relative weights could differ across populations.

Polyploidy is relatively frequent in *S. cerevisiae* and, as found in this study, strongly associated with heterothallism. Since mating-type switching bypasses incompatibility between haploid cells, the primary role of mating types in *S. cerevisiae* may not be to prevent selfing, and the cost of inbreeding, as in most outbreeding organisms. Instead, the “developmental switch model” has suggested that mating types function as a ploidy sensor (2, 29, 30): *MATa* and *MAT*α cells express haploid-specific genes, which prevent meiosis and allow mating, whereas *MATa*/*MAT*α cells express diploid-specific genes, which prevent mating and allow meiosis. Thus, mating types ensure the alternation between haploid and diploid phases by preventing aberrant meiosis or mating. We propose that heterothallism might promote polyploidy by extending the duration of haploid phases during vegetative growth. In homothallic lineages, *MATa* and *MAT*α cells convert to *MATa*/*MAT*α within at most two mitotic divisions, making haploid states both rare and less exposed to selection than the diploid state. By contrast, heterothallic strains likely spend more time and undergo more divisions in the haploid states. This state is more prone to whole-genome endoreduplication (24, 31, 32), which generates diploid cells with a single mating type. These cells would be able to mate with a cell of the opposite mating type, leading to the formation of a polyploid. In an alternative scenario, polyploidization could occur first and make the use of selfing unnecessary, either because polyploids can produce spores carrying the two mating types, or because sexual cycles would rarefy, which would relax the selective pressure on the *HO* gene.

Altogether, our findings show that the evolution of mating-type switching in *S. cerevisiae* is highly dynamic, with repeated transitions to heterothallism reshaping life cycle strategies. These changes profoundly affect genomic diversity by influencing patterns of heterozygosity and polyploidy. The causal order between heterothallism, heterozygosity and polyploidy remains to be fully demonstrated. The high proportion of heterozygous heterothallic isolates (91%) supports a model in which heterothallism promotes heterozygosity by preventing complete homozygotization through haploid selfing. The causal relationship with polyploidy is less clear, but the hypothesis that heterothallism promote polyploidy by prolonging haploid vegetative growth is directly testable. More broadly, our results demonstrate that repeated evolutionary changes in a single life-history trait can shape genome architecture and drive ecological diversification within a species.

## Materials and Methods

### Strain panels and genome

The *Saccharomyces cerevisiae* Reference Assembly Panel (ScRAP, (17)) contains 140 strains and their telomere-to-telomere (T2T) genome assemblies (Dataset S1). The S288C and CGH_3 assemblies were excluded because they were redundant with the SGDref and CGH_1 assemblies, respectively. BLAST databases were created using makeblastdb from blast v. 2.15.0 (33). Identification of artificial gene disruptions was done with blastn from blast 2.15.0 (33), using the sequences of the *Ashbya gossypii* TEF promoter (NCBI id: MK178574.1:6908-7251) and terminator (MK178574.1:8067-8264), Hyg1 (EF101286.1:459- 1487), Kan1 (KJ502278.1:1963-2772) and Nat1 (MK431404.1:801-1370). Hits in chromosome IV were considered as artificial disruptions of the *HO* gene, and the 23 corresponding assemblies and strains were discarded from further analyses (Dataset S2). For the 117 remaining strains, the ScRAP’s phylogenetic tree and the genome assemblies’ statistics were retrieved from (17). Strains were considered as monosporic isolates when annotated as “monosporic” and “*de novo* sequenced and assembled” in the Table S1 of (17). The ploidy of each isolate were retrieved from the Table S1 of (17), except for 9 isolates for which we found inconsistencies (AAB, ACA, AEH, AFH, AFI, AGK, CDA, HLJ1, SK1): ploidy of AAB, ACA, CDA and AFI was then inferred by flow cytometry (AAB and AFI were diploid while CDA and ACA were haploid), and remained unknown for the others. For the nine isolates, zygosity was set to unknown.

The 3,034 Genome Panel (3,034GP (28)) contains 3,039 strains. Metadata were downloaded from the Table S1 of (28). We used supplementary data from (13) to exclude 124 strains with an indicated artificial *HO* deletion.

### Media and ascus dissection

Cells were grown at 30°C in liquid YPD (5% YPD Broth, Sigma-Aldrich) or on solid YPD (6.5% YPD Agar, Sigma-Aldrich). Cells were sporulated for 7 days on solid sporulation media (2% Potassium Acetate and 2.5% BactoAgar, Sigma-Aldrich), the presence of asci was checked by optical microscopy. Asci were incubated in 10 μL H_2_O with 100 μg/mL of lyticase at 37°C for 6 minutes to digest the ascus wall, then dissected with a micromanipulator (Singer MSM System 400). Monosporic isolates, derived from a single haploid spore, were recovered after three days of growth at 30°C on YPD plates.

### *MAT* genotyping by polymerase chain reaction (PCR)

The *MAT* locus was genotyped by PCR using one common forward primer outside of the *MAT* locus (CAAGGGAGAGAAGACTTGTG), and one reverse primer specific for MATa (CCACTTCAAGTAAGAGTTTGG) and MATα (AGCACGGAATATGGGACTAC). As a positive PCR control, we used two primers (GTATGATGAAAGCCAATTCACC and CGGTATGTCTCGAGTATTACC) to amplify a sequence within the gene *TAF2*. Cells from 1 to 3 days growth on solid YPD were placed in H_2_O at -20°C overnight, then at 95°C for 10 minutes, then on ice for 10 minutes, then centrifugated for 30 seconds to pellet cellular debris. 10 µL supernatant was used as matrix for the PCR reaction in a final volume of 50 μL containing 1.25 U of DreamTAQ DNA Polymerase (Thermo Scientific), 5 μL of 10X DreamTaq Buffer (Thermo Scientific), 0.5 μM of each of the five primers and 200 μM of each dNTP. The PCR program contained a 2-minute initial denaturation at 95°C, then 35 cycles (95°C for 30 seconds, 55°C for 30 seconds, 72°C for 1 minute) and a 10-minute final extension at 72°C. Amplicon visualization was made by electrophoresis on 1%-agarose gels with 10μL PCR product per well. Three control isolates were used for the PCR: BY4741 (MATa) and BY4742 (MATα) and the diploid strain BY4743 (MATa/MATα).

### Molecular complementation assay

The pHS2 plasmid (gift from John McCusker, RRID:Addgene_81037) contains a functional *HO* locus (the 1761bp coding sequence, flanked by the 766bp upstream and 823bp downstream) as well as a HYGMX locus conferring resistance to hygromycin. For transformation, overnight yeast culture was diluted 100 times in liquid YPD and incubated 4 hours at 30°C. Then 10^8^ cells were washed twice in sterile distilled water, then washed three times in 0.1M Lithium Acetate Tris Ethylenediaminetetraacetique (LiAcTE). The cells were then incubated at 42°C for 25 minutes with 30 μL 1M LiAcTE, 270 μL of a 50% solution of Propylene Glycol, 50μg of single-strand salmon sperm DNA and 500 ng of pHS2 plasmid. Then, cells were incubated in liquid YPD for 2 hours to allow the expression of the hygromycin resistance genes. Cells were then plated on solid YPD + 200 µg/mL hygromycin. Two independent transformant clones (or one when there was one colony on the selection plate) were picked from each transformation experiment. For each clone, the *MAT* locus was genotyped by PCR.

### Ploidy estimation by flow cytometry

A total of 5.10^6^ cells of an overnight culture in YPD were incubated at 4°C overnight in 70% ethanol, then washed twice in PBS, resuspended in 500μL of staining solution (15 μM Propidium Iodide, 100 μg/ml RNase A, 0.1% v/v Triton-X, in PBS), then incubated for 3 hours at 37°C in darkness and sonicated for 15 seconds. A sample of 10000 cells was analyzed with a MACSQuant VYB Flow Cytometer. Debris and multiplets were removed from the analyses as described above. Cells were excited at 561 nm and fluorescence was collected with a 615/20 filter. Data analyses were performed with the R package CytoExploreR v1.1 [3]. The values of the peaks corresponding to G1 and G2 phases relative to the control strains were used to indicate the ploidy of each sample. Three control isolates were used: BY4741 (haploid), BY4742 (haploid) and BY4743 (diploid).

### Analysis of the *HML***α** and *HMRa* repair cassettes in ScRAP

The sequences of X, Ya, Yα and Z1 regions of the *MATa*/*HMRa* and *MAT*α/*HML*α, as well as two surrounding genes per locus (*VBA3* and *CHA1* for *HML*, *PHO87* and *TAF2* for *MAT*, *CDC50* and *GIT1* for *HMR*) were downloaded from the SGD website and blasted on the ScRAP nuclear assemblies using the blastn command from blast 2.15.0 (33). For heterozygous strains, we retained hits from phased assembly only. We kept only the hits that mapped between the pairs of surrounding genes for each locus.

### Functional analysis of the *HO* coding sequence in ScRAP

*HO* coding sequences were identified in the ScRAP assemblies using the blastn command from blast 2.15.0 (33), with the *HO* sequence from plasmid pHS2 as the query. For heterozygous isolates, we retained hits from the phased assembly when available. If no hit was found in the phased assembly, the hit from the collapsed (haploid consensus) assembly was used instead. Partial hits located at scaffold extremities and one low-quality hit on block100_contig2 of the ANL assembly were excluded. Contiguous hits on the same scaffold were merged, as they likely reflected deletions rather than distinct loci. The resulting sequences were extracted using the getfasta command of bedtools v2.31.1 (34) and aligned with MAFFT v7.526 (35). Variants relative to the pHS2 reference were identified and annotated by type (single nucleotide polymorphism [SNV], insertion, deletion). Variants were classified into three categories based on their predicted impact on Ho protein function, taking into consideration (i) their impact on the coding sequence, (ii) the results from the experimental screen, (iii) the results from the complementation assay and (iv) published data:

1. Neutral: synonymous variants, variants found in assemblies from homothallic isolates and variants previously described as neutral in (23).
2. LoF: the 667G>A variant demonstrated to inactivate *HO* in (23), non-synonymous variants at functionally critical positions described in (27) and variants resulting in nonsense mutations, start codon loss, or frameshifts.
3. Unknown: all other variants.

The *HO* alleles are unique combinations of variants present in the MSA. They were classified into three functional categories based on their variants, and experimental evidence:

A. Functional alleles: contain only neutral variants.
B. Non-functional alleles: contain at least one LoF variant, or present in the assembly of a heterothallic isolate which rescued switching in complementation assays.
C. Unknown alleles: all remaining cases.

The thallism of the isolates that were not tested during the experimental screen was predicted as follows: if all alleles present in a genome assembly are functional, the isolate is homothallic and if all alleles present in its genome assembly are non-functional, the isolate is heterothallic. We found no strain with both functional and non-functional alleles of *HO*.

### Computing the minimal number of HO loss-of-function events in ScRAP

We looked for ***S***, the smallest set(s) of non-neutral variants (categories 2 and 3) so that all non-functional alleles (category B) carry at least one variant. The cardinal of ***S*** is denoted ***M***: it corresponds to the minimal number of non-neutral variants required to explain that all the alleles are non-functional. ***M*** estimates the minimal number of independent events triggering a novel loss of function of the protein, since additional LoF mutations could have accumulated once the gene is already non-functional. We computed ***M*** as follows:

1. First, for each allele with a single variant, this variant is necessarily included in ***S***. This represents ***M_1_*** variants. All alleles carrying these variants were ignored for the next step.
2. Then, on the remaining alleles, a greedy algorithm based on selecting recursively the most frequent variant was used to obtain a first suboptimal solution ***M_G_***.
3. To find the optimal solution, at least one solution was searched in all sets of size ***M_G_ – 1***, then of size ***M_G_ – 2***, …, until no solution is found. The last successful size is denoted ***M_2_***.
4. Finally, we could compute the size of ***S***: ***M*** *= **M_1_** + **M_2_***.

### *HO* sequence network

*HO* sequence network was built using SplitsTree CE 6.0.0 (36), with as input the multiple sequence alignment of the 151 *HO* sequences from ScRAP. The P Distance method (37) was used so as to obtain a 151 x 151 distance matrix. The Neighbor Net method (38, 39) was used so as to obtain 243 splits, cyclic. The Show Splits method was used so as to obtain a Split Network visualization.

### Predicting the thallism of the 3,034GP

The VCF file of the *HO* locus (chromosome 4, positions 46271 to 48031) from the 3,034GP was provided by Jing Hou ((28)) and is available on the following Zenodo repository: https://zenodo.org/records/17106281. One *HO* sequence (with IUPAC code for heterozygous sites) per strain was reconstructed using the consensus command from bcftools v. 1.20 (40) and the reference genome from the Zenodo repository of (28) https://doi.org/10.5281/zenodo.12580561. Sequences from the 124 strains with an indicated artificial *HO* deletion in (13) were excluded. The 2,915 remaining sequences were aligned with MAFFT v7.526 (35). Variants relative to the pHS2 functional reference were identified and annotated by type (single nucleotide polymorphism [SNV], insertion, deletion), predicted effect on the coding sequence (synonymous, missense, nonsense, start codon loss, in-frame insertion or deletion, or frameshift), and presence/absence across sequences in the alignment. We found no variant between positions 1560 and 1681, which is the position of the 108bp in-frame deletion that we identified in the ScRAP sequences. Variants were classified into three categories based on (i) their predicted impact on Ho protein function, considering their coding effect, (ii) results from the ScRAP *HO* haplotypes, and (iii) published data:

1. Neutral: synonymous variants and neutral variant identified in the ScRAP analyses of *HO* haplotypes.
2. LoF: the 667G>A variant identified to inactivate *HO* in (23), non-synonymous variants at functionally critical positions described in (27), and variants resulting in nonsense mutations, start codon loss, or frameshifts.
3. Unknown: all other variants.

A strain was considered as homothallic if its *HO* sequence contained only neutral variants. It was considered as heterothallic if it carried at least one LoF variant: fully heterothallic if the LoF is homozygous in this genotype and partly heterothallic if the LoF is heterozygous.

### Classification of ecological niches

The ecological origin of each strain origin was classified as “Anthropogenic”, “Natural” or “Unknown”, based on the presence of keywords in the ‘EcoOrigin column of Table S1 from (28). The keywords associated with the “Anthropogenic” niche are: active, alcohol, ale, apple, avocado, baker, baking, banana, beer, belgian, bioethanol, biofuel, blood, bread, brewery, burukutu, cacao, cachaca, cake, cane, champagne, cheese, cider, citrus, clinical, cocoa, coconut, coffee, commercial, corn, crop, cultivated, culture, dairy, derivative, dietary, dietetic, distillery, distilling, dough, dried, feces, feed, ferment, figs, flor, fluid, food, fruit, fuel, grape, guava, hibius, human, industrial, kefir, kefyr, kumiss, kunnu, lab, lager, lungs, lychee, man, market, mash, milk, orange, orchard, papaya, patient, peache, potato, probiotic, rhum, sake, sap, sherry, slope, spirits, sputum, starch, starter, tapioca, togwa, tokay, vagina, vineyard, wheat, white tecc,wine. The keywords associated with the “Natural” niche are: animal, bark, beetle, butterfly, canyon, cassine, castanea, drosophila, fagus, ficus, fish, flower, forest, fraxinus, insect, leaf, leave, mushroom, nature, oak, plant, pseudoobscura, pyrenaica, quercus, soil, symbiosis, tapirira, tree, water, wild. When keywords from both lists are found in the same niche description, the ecological niche is classified as “Anthropogenic”. If no keyword was found, the niche was classified as “Unknown”.

### Data, Materials, and Software Availability

Data and codes generating the figures and tables are deposited the following Zenodo repository: https://zenodo.org/records/17106281. All other data are included in the manuscript and/or in the supporting

## Acknowledgments and Funding Sources

We are grateful to our colleagues in the Biology of Genomes, Stochastic Models for the Inference of Life Evolution and Telomere Stability groups for their valuable discussions and insights, and to Tamera Lahrache for technical support. This work was supported by the Agence Nationale de la Recherche ANR-20-CE12-0020.

## Notes

https://zenodo.org/records/17106281

### Competing Interest Statement

The authors have declared no competing interest.

