## Supporting Information for "Repeated losses of self-fertility shaped heterozygosity and polyploidy in yeast evolution"

**This PDF file includes:**

Supporting Information text

Figures S1 to S7

Legends for Datasets S1 to S5

SI References

**Other supporting materials for this manuscript include the following:**

Datasets S1 to S5

Supporting Information Text

The *Saccharomyces cerevisiae* Reference Assembly Panel

The *Saccharomyces cerevisiae* Reference Assembly Panel (ScRAP, (1)) contains 140 strains and their telomere-to-telomere (T2T) genome assemblies. Genomes assemblies were downloaded from <https://www.evomicslab.org/db/ScRAPdb/download/>. The S288C and CGH_3 assemblies were excluded because they were redundant with the SGDref and CGH_1 assemblies, respectively. BLAST databases were created using makeblastdb from blast v. 2.15.0 (2). Identification of artificial gene disruptions was done with blastn from blast 2.15.0 (2), using the sequences of the *Ashbya gossypii* TEF promoter (NCBI id: MK178574.1:6908-7251) and terminator (MK178574.1:8067-8264), Hyg1 (EF101286.1:459-1487), Kan1 (KJ502278.1:1963-2772) and Nat1 (MK431404.1:801-1370). Hits in chromosome IV were considered as artificial disruptions of the *HO* gene in 23 assemblies and the corresponding strains were discarded from further analyses (Dataset S1). In two isolates (CBK and CBM), the markers were detected elsewhere in the genome, but not within the *HO* locus. As there is no evidence suggesting any artificial modification of their *HO* sequences, these two isolates were retained for further analyses. For the 117 strains with no artificial *HO* deletion, the ScRAP phylogenetic tree and the genome assembly statistics were downloaded from <https://www.nature.com/articles/s41588-023-01459-y#Sec40>. Isolates were considered as monosporic isolates only when annotated as “monosporic” and “*de novo* sequenced and assembled” in the Table S1 of the ScRAP paper. The ploidy and zygosity of each isolate were retrieved from the Table S1 of the ScRAP paper, except for nine isolates for which the ploidy and zygosity were settle to “unknown”, including seven isolates which were annotated as “ho::KanMX” in the metadata but showed no evidence of artificial disruption of their *HO* gene in the assembly (AAB, ACA, AEH, AFH, AFI, AGK, CDA), and two isolates with inconsistent ploidies between Tables S1 and S2 (HLJ1, SK1). We inferred the ploidy of AAB, ACA, CDA and AFI by flow cytometry and found that AAB and AFI were diploid while CDA and ACA were haploid. The ploidy of the other five strains remained undetermined. For the nine isolates, zygosity was set to unknown. The ploidy and zygosity of the original strains that were used to derive the monosporic isolates were retrieved from Table S2 of (1).

Experimental screen for homothallic and heterothallic strains in ScRAP

Among the 140 isolates from the ScRAP (1), 23 had an artificial *HO* deletion. Among the 117 remaining strains, 7 were absent or misidentified in our stock (CENPK1137D, SK1, SX2, Y55, ADS, AVB, AEG). Among the 110 remaining strains, we excluded 13 polyploids and the 4 isolates with uncertain ploidy, leaving 93 strains for the experimental screen, 45 of which being monosporic isolates derived from a single spore of an originally diploid strain and 48 being original isolates.

We first tested the 45 ScRAP monosporic isolates. We genotyped their *MAT* locus by PCR. Three control isolates were used for the experiments: BY4741 (MATa) and BY4742 (MATα) and the diploid strain BY4743 (MATa/MATα). A total of 13 isolates with genotype *MATa* or *MATα* were considered as heterothallic. *MATa/MATα* diploids were considered as homothallic (n = 29). One diploid monosporic isolate (AMH_1a) was contaminated in the stock, and its PCR genotyping failed. We dissected new monosporic isolates from its parent (AMH), and found that all carried the *MATa/MATα* genotype, confirming that AMH_1a was also a homothallic *MATa/MATα* diploid. Two monosporic isolates (CHS_3a and CPI_1c) were annotated as haploid in the original paper, but carried *MATa/MATα* genotype, possibly resulting from rare homologous recombination at the *MAT* locus (3) and not classical mating-type switching. We dissected new monosporic isolates from these two genetic backgrounds, and found that all carried a single *MATa* or *MATα* allele and were unable to sporulate, confirming that these two isolates are actually heterothallic. For these two isolates, we looked at the *MAT* genotype in the genome assembly and found that they both carry the *MATa* allele. Since they probably became *MATa/MATα* after sequencing, we considered that these strains were haploid *MATa* for our analyses.

We then genotyped the *MAT* locus of the 48 original strains by PCR. We identified 13 isolates with *MATa* or *MATα* genotypes, which we considered heterothallic. We sporulated the 35 remaining *MATa/MATα* diploids and found that 14 had impaired sporulation (10 no sporulation at all, 3 very low sporulation rate and 1 produced only dyad) and two produced no viable spores. For the 19 remaining strains, 5-30 tetrads were dissected on solid YPD, *i.e*., each spore of the tetrad was deposited at a determined spot of the plate. Spore survival was determined by the presence of a colony at the spot after 3 days of growth. For each isolate with non-zero spore survival, 6-30 new monosporic isolates were genotyped for their *MAT* locus. When less than 10% (or less than one) new monosporic isolates were MATa/MATα, the isolate was considered heterothallic. When less than 10% (or less than one) new monosporic isolates were *MATa* or *MATα*, the isolate was considered homothallic.

Analysis of the *MAT*, *HML* and *HMR* loci in the ScRAP

The *HML*, *HMR* and *MAT* loci are all composed of three adjacent regions, X, Y and Z1. X and Z1 have similar sequences in the three loci. They flank the Y region, which has two alleles, Ya (in *MATa* and *HMRa*) and Yα (in *MATα* and *HMLα*). The sequences of X, Ya, Yα and Z1, as well as the sequences of two surrounding genes per locus (*VBA3*-YCL069W and *CHA1*-YCL064C for *HML*, *PHO87*-YCR037C and *TAF2*-YCR042C for *MAT*, *CDC50*-YCR094W and *GIT1*-YCR098C for *HMR*) were downloaded from the SGD website and blasted in the ScRAP nuclear assemblies using the blastn command from blast 2.15.0 (2). For each heterozygous strain, we retained hits from its phased assembly only. Because of the repetitive nature of the sequences in *HML*, *HMR* and *MAT*, we kept only hits between a pair of surrounding genes for each locus (VBA3 and CHA1 for *HML*, PHO87 and TAF2 for *MAT*, CDC50 and GIT1 for *HMR*). A X-Y-Z1 sequence was considered as a *HML* locus when it was present between hits for its surrounding genes VBA3 and CHA1, as a *HMR* locus when it was present between hits for its surrounding genes CDC50 and GIT1.

We found a total of 111 *HMR* loci from the genome assembly of 100/117 strains, all contained the X-Ya-Z1 sequence as expected. The *HMR* locus from BJ4 exhibits a partial deletion of the Ya region and the full Z1 sequence is lacking. We retrieved 121 *HML* loci from the genome assembly of 102/117 strains, among which 109 contained the X-Yα-Z1 sequence as expected, and 12 contained an unexpected X-Ya-Z1 sequence. Missing loci may be due to low contiguity in some genome assemblies and not only to real absence.

*HO* sequences in the ScRAP

In 85 haploid assemblies, comprising 45 monosporic isolates, 9 haploids, 21 homozygous diploids, 1 homozygous triploid, and 9 strains with unknown ploidy, we identified a single *HO* sequence per assembly, for a total of 85 sequences. In the 20 phased assemblies from heterozygous diploids, 38 *HO* sequences were recovered, with two sequences per assembly except for CFF and CBM, where only one complete sequence was found due to the second being truncated at a scaffold end. In the phased assemblies of the 5 triploid isolates, we identified a total of 6 *HO* sequences: 3 in BBT as expected, 2 in CPS, and one in CRE, but no sequences were recovered in BAD or CGH from the phased assemblies. In the seven phased tetraploid isolates, nine *HO* sequences were found: four in ANL, ANM, and BTE as expected, three in ATV, two in AVN and BEM, and none in CFC. For the 3 polyploid isolates where no *HO* sequence could be retrieved from the phased assemblies (BAD, CGH, and CFC), one sequence per isolate was extracted from their collapsed (*i.e.*, haploid consensus) assembly. The number of *HO* sequences recovered in some diploid and polyploid isolates was lower than expected based on their ploidy, likely due to technical limitations in genome assembly and phasing rather than the true absence of the gene.

Predicted *HO* functionality and homo- or heterothallism in the ScRAP strains

We identified 80 different *HO* alleles (*i.e.*, unique combinations of variants) among the 151 *HO* sequences. We considered that an allele is functional when it carries only neutral variants. We found 32/80 functional alleles, corresponding to 63/151 sequences and present in 59/117 isolates (Supplementary Figure 4). Conversely, we considered that an allele is non-functional when it carries at least one LoF variant or if it was characterized as such in the complementation assay. We found 33/80 non-functional alleles, corresponding to 65/151 sequences and present in 43/117 isolates (Supplementary Figure 4). We also used the predicted functionality of *HO* alleles to infer the switching phenotype of the 40 ScRAP isolates that could not be tested experimentally (Dataset S6). Predictions were based solely on *HO* allele functionality, which is a reliable proxy for switching ability in single-mating-type contexts. Some of these strains, however, are polyploid. If they sporulate, they can produce spores with diverse MAT genotypes, and only those with a single MAT genotype are formally relevant for defining homo- or heterothallism. For clarity, we nonetheless describe the polyploid strains themselves as homo- or heterothallic. Among the 40 untested strain, 16 carried only non-functional *HO* alleles and are therefore predicted to be heterothallic, 14 carried only functional *HO* alleles and are predicted to be homothallic, and no prediction could be made for the remaining 10 strains. Overall, we estimated that 48% (57/117) of ScRAP isolates are homothallic, 41% (48/117) are heterothallic, 2% (2/117) exhibit both phenotypes, and 8% (10/117) have an unknown phenotype.

We looked at the predicted functionality of the *HO* alleles in the heterothallic strains that show no complementation upon transformation with a functional copy of *HO*. Only one (ADM) carried an allele of unknown functionality, whereas the other four (AAC, AEL, AQG_2a, CDA) carried predicted non-functional *HO* alleles. Among the nine heterothallic isolates that were not tested in the complementation assay, one carried a functional allele (AAB), three carried alleles of unknown functionality (ADQ, BGP_1a, CPI_1c) and five (ACA, CBK, CBM, CHS_3a, S228c) carried predicted non-functional *HO* alleles. These results further support the conclusion that loss-of-function mutations in *HO* are the main cause of heterothallism.

The 3,034 Genome’s Panel (3,034GP (4))

The 3,034 Genome Panel (3,034GP (4)) contains 3,039 strains. Metadata, including ploidy and genome-wide zygosity were downloaded from the Table S1 of (4). Strains containing more than 500 heterozygous SNPs were considered heterozygous whereas the rest were considered homozygous. Ploidy was estimated by flow cytometry in only a fraction of the collection (1,278 strains) whereas for the other strains, ploidy could be inferred only in heterozygous strains based on their allele balance ratio. This is a bias to consider when looking at association between ploidy, genome-wide zygosity and, in our case, thallism. We used supplementary data from (5) to exclude 124 strains with an indicated artificial *HO* deletion, keeping 2,915 strains for the rest of the study (Supplementary Figure 6).

*HO* genotypes in the 3,034GP (4)

The VCF file of the *HO* locus (chromosome 4, positions 46271 to 48031) from the 3,034 Genome’s Panel was provided by Jing Hou ((4), Supplementary Files). One *HO* sequence (with IUPAC code for heterozygous sites) per strain was reconstructed using the consensus command from bcftools v. 1.20 (6) and the reference genome from the Zenodo repository of (4) <https://doi.org/10.5281/zenodo.12580561>. Sequences from the 124 strains with an indicated artificial *HO* deletion in (5) were excluded. The 2,915 remaining sequences were aligned with MAFFT v7.526 (7). Variants relative to the pHS2 functional reference were identified and annotated by type (single nucleotide polymorphism [SNV], insertion, deletion), predicted effect on the coding sequence (synonymous, missense, nonsense, start codon loss, in-frame insertion or deletion, or frameshift), and presence/absence across sequences in the alignment. We found no variant between positions 1560 and 1681, which is the position of the 108bp in-frame deletion that we identified in the ScRAP sequences. Since this variant was rather frequent in the ScRAP, it may also have been frequent in the 3,034GP and have prevented variant calling in this region. We did not find variants between positions 1708 to 1761 either. This suggests that we probably missed variants in these regions, leading to underestimate the frequency of heterothallic isolates (only neutral variants) and to overestimate the frequency of homothallic (no LoF variant). We found 395 variants, among which 247 where present in a heterozygous state in at least one strain (63% of the variants). Among the 2915 strains, 796 (27%) had at least one heterozygous site in *HO*. Among the 395 variants, we found 127 synonymous SNVs, 239 missense SNVs, 11 nonsense SNVs, 16 indels causing a frameshift, one in-frame insertion and one loss of start codon. We considered that 40 variants were LoF, including the 11 nonsense SNPs, the 16 frameshift, the loss of start codon, the 667G>A variant demonstrated to inactivate *HO* in (8) and 11 missense SNVs at the functional critical positions described in (9) – 646C>A, 646C>T, 649G>C, 662G>A and 668G>A in the first LAGLIDADG motif, 983G>C and 997G>A in the second LAGLIDADG motif, 1405T>A in the first finger of the 1^st^ zinc finger, 1522T>C and 1531T>A in the first finger of the 2^nd^ zinc finger, and 1682G>A in the first finger of the 3^rd^ zinc finger.

Figures

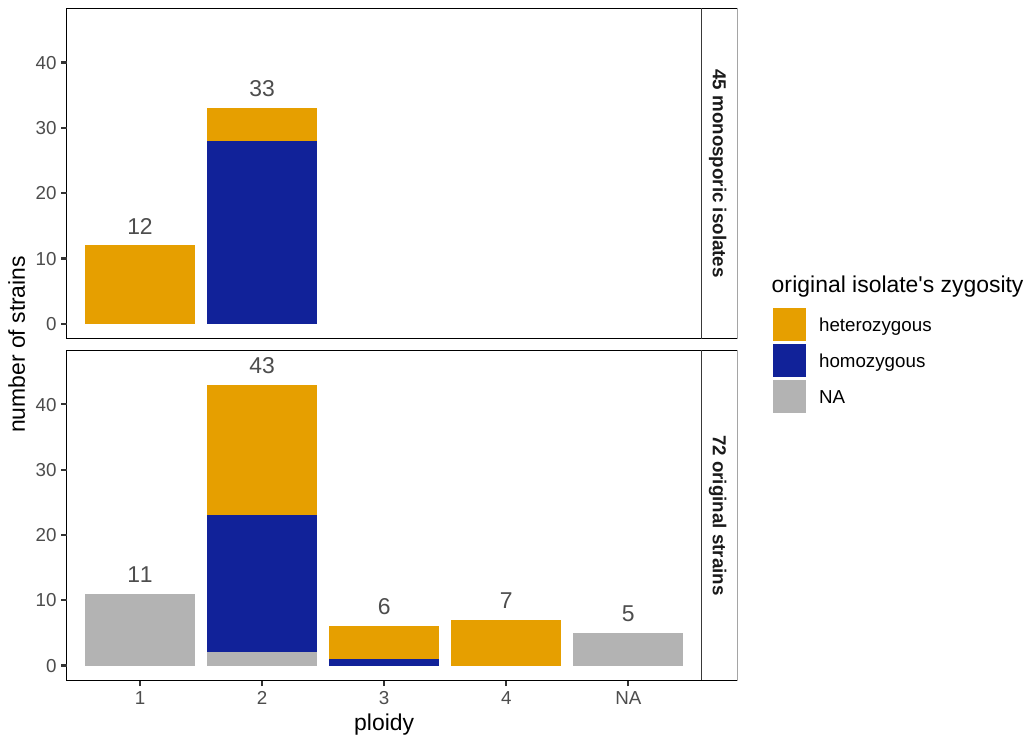

Fig. S1. Distribution of ploidy and zygosity across the ScRAP. Among the 140 strains, 23 were excluded because they have an artificial *HO* deletion. The remaining 117 strains comprises 45 monosporic isolates derived from a single spore of an originally diploid strain (top), and 72 original strains (bottom). Color represent the genome-wide zygosity, which was defined in (5): strains with less than 5% of heterozygous sites were considered as homozygous.

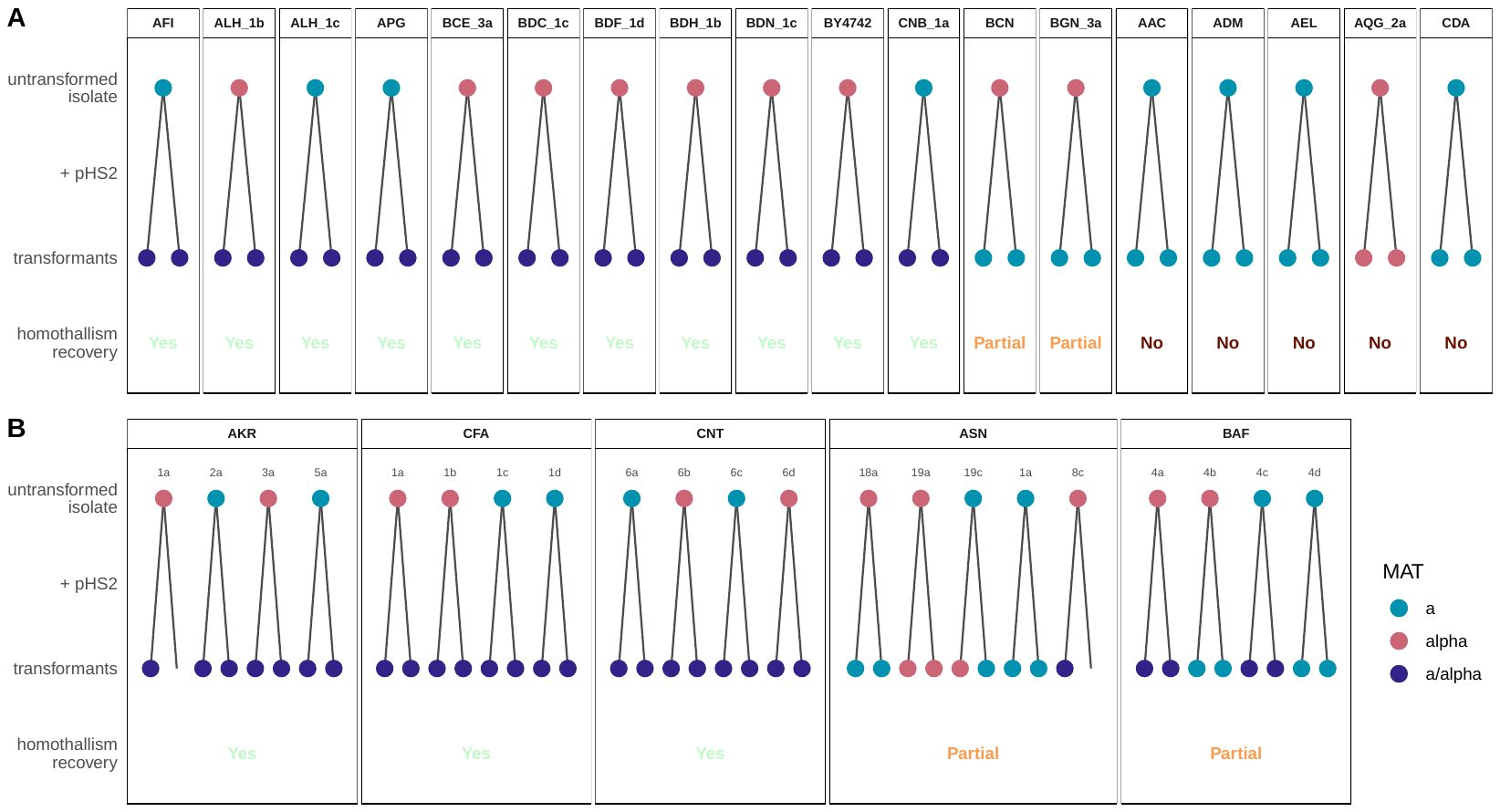

Fig. S2. Complementation of heterothallic strains with a functional *HO* gene. (A) Single-mating type isolates. (B) Diploid *MATa*/*MATα* isolates: multiple monosporic isolates were tested – their id is indicated in grey, with a number for the tetrad and a letter for the spore. The dots represent the isolates before (top) and after (bottom, two replicates) transformation with the pHS2 plasmid containing a functional *HO* gene; colors correspond to the *MAT* genotype. A single transformant was obtained from the transformations of AKR.1a and ASN.8c. The four monosporic isolates from the same tetrad were tested when spore viability allowed it.

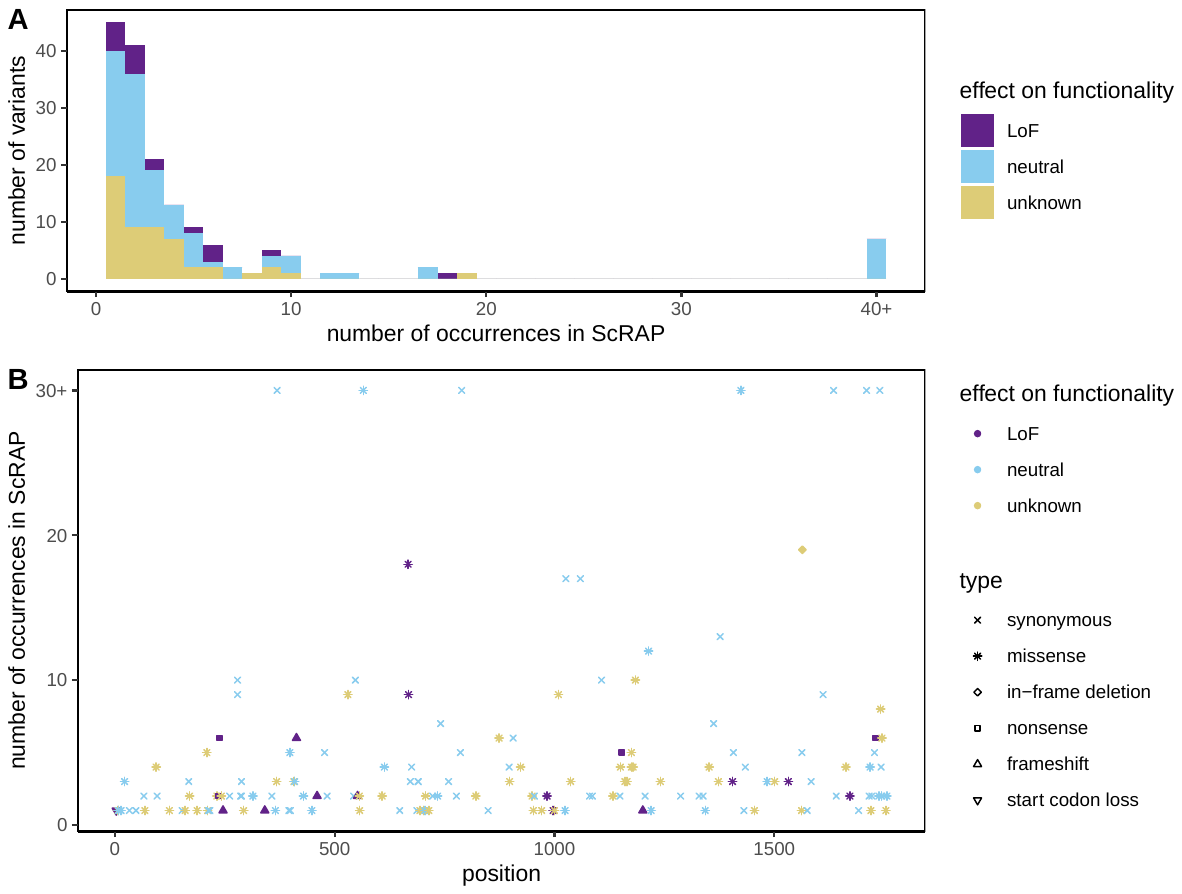

Fig. S3. Variants present in the *HO* coding sequence of ScRAP. (A) Number of occurrences of the variants in the 151 coding sequences of ScRAP. (B) Position of the variants on the 1,761 bp coding sequence of *HO*.

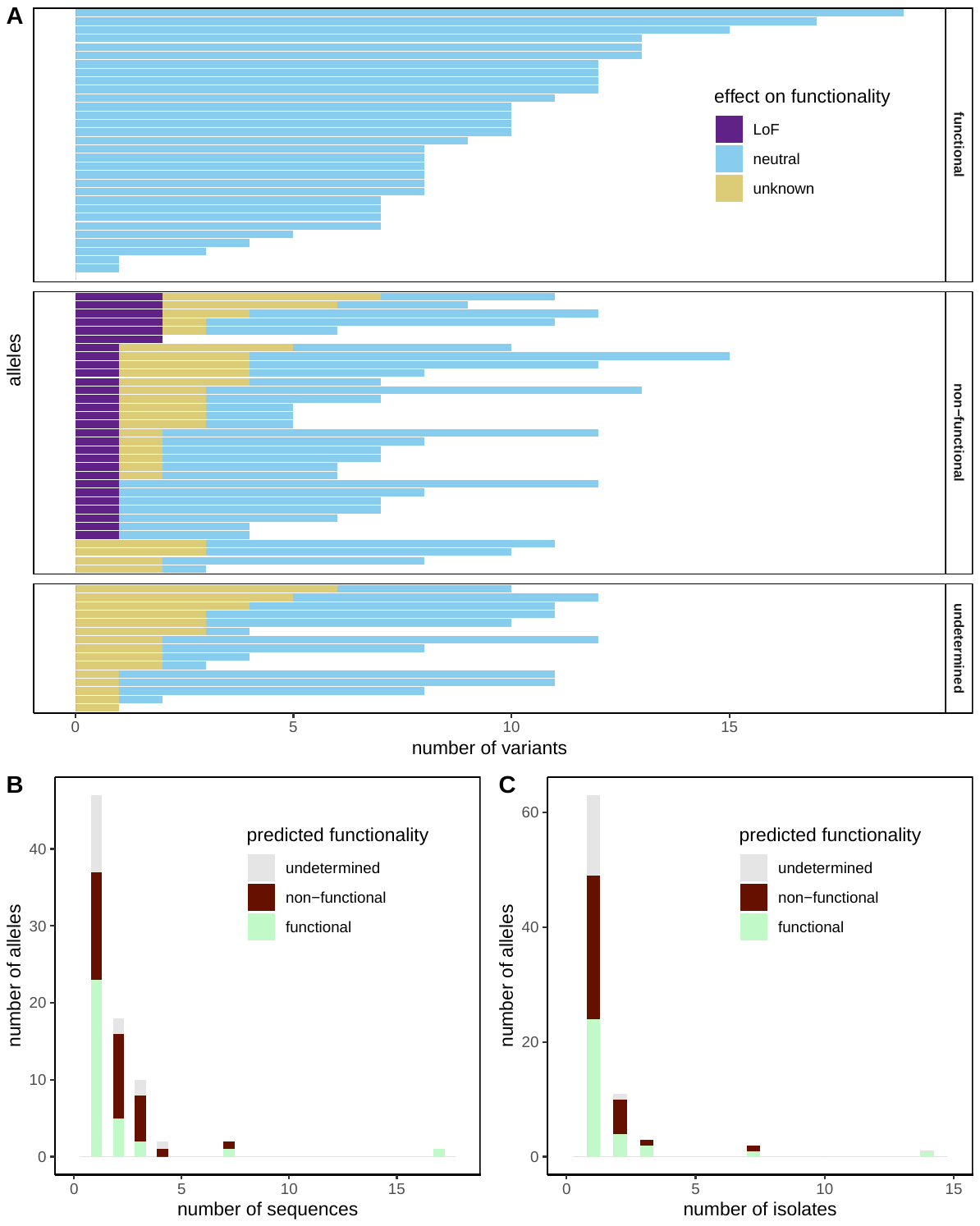

Fig. S4. Predicted functionality of *HO* alleles in the ScRAP. Allele functionality prediction was based on the type of variants present in the allele and the experimental results of this study. (A) Number and effect of the variants carried by the 80 *HO* alleles. (B) Distribution of the alleles in the 151 coding sequences of *HO*. (C) Distribution of the alleles in the 117 isolates’ genome assemblies.

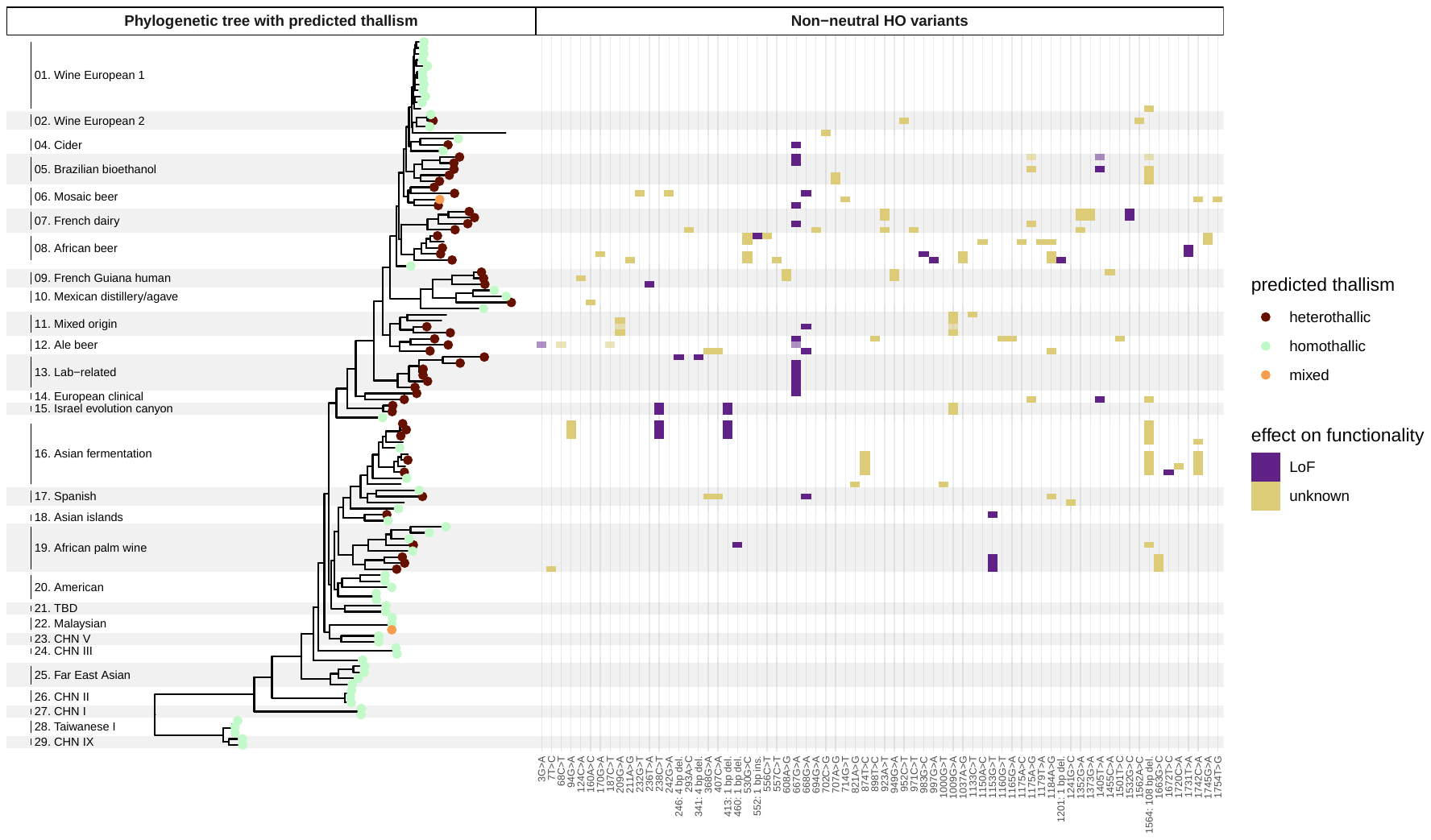

Fig. S5. Phylogenetic distribution of homo- and heterothallic strains and non-neutral variants from the ScRAP. Left: Predicted thallism of ScRAP strains, mapped on the phylogenetic tree from the whole-genome date from (1). The names on the left and the grey and white rectangles stripes the different clades identified in (1). Leave colors represent predicted thallism, based on experimental data from this study and predicted functionality of the *HO* allele. Right: Presence of the 70 non-neutral *HO* variants – comprising the 18 LoF variants and the 52 variants with unknown effect. Lighter square indicates that the variant is present at the heterozygous state.

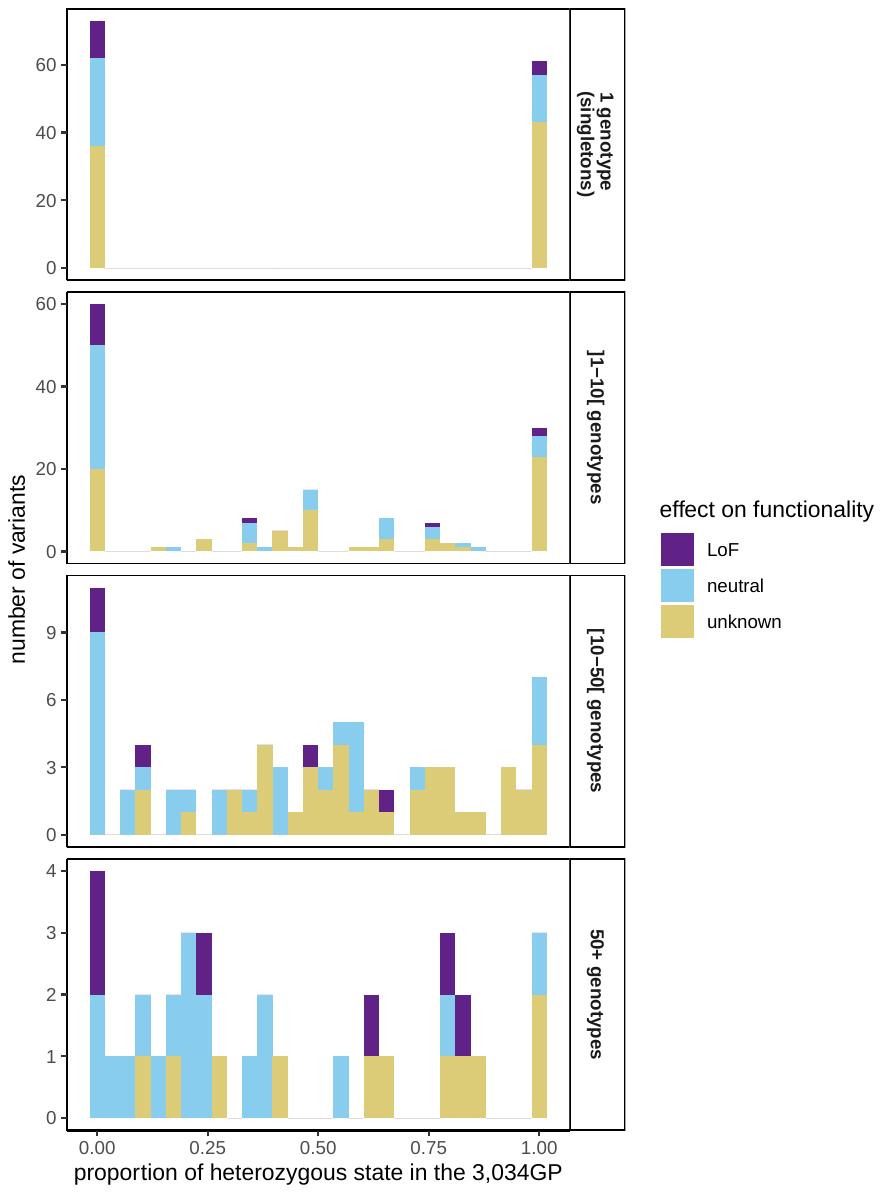

Fig. S6. Proportion of heterozygous state of the 395 *HO* coding sequence variants in the3,034GP. On the y-axis, the variants were classified according to the number of genotypes in which they are found.

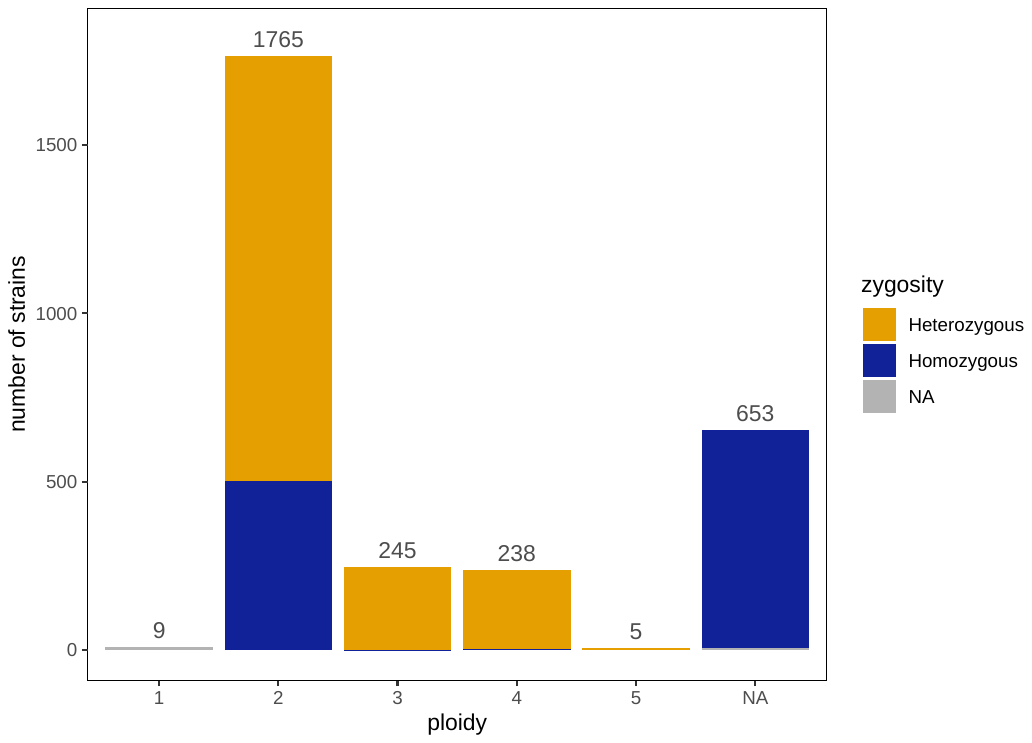

Fig. S7. Distribution of ploidy and zygosity across the 3,034GP. Among the 3,039 strains, 124 were removed because of an artificial deletion. Here are represented only the 2,915 strains that passed this filter. We retrieved genome-wide zygosity and ploidy information for each strain from the original paper of the 3,034GP (4). Strains containing more than 500 heterozygous SNPs were considered heterozygous whereas the rest were considered homozygous. Ploidy was estimated by flow cytometry in only a fraction of the collection (1,278 strains) whereas for the other strains, ploidy could be inferred only in heterozygous strains based on their allele balance ratio.

**Dataset S1 (separate file).** The 117 strains from the ScRAP analyzed in this study.

**Dataset S2 (separate file).** Markers of artificial gene deletions in the detected in the ScRAP.

**Dataset S3 (separate file).** *HO* coding sequence variants in the ScRAP.

**Dataset S4 (separate file).** *HO* coding sequence variants in the 3,034GP.

**Dataset S5 (separate file).** The 2915 strains from the 3,034GP analyzed in this study.
